# Cell-mediated cholesterol crystal processing and clearance observed by 3D cryo-imaging in human atherosclerotic plaques

**DOI:** 10.1101/2023.11.28.568890

**Authors:** Jenny Capua-Shenkar, Antonia Kaestner, Katya Rechav, Vlad Brumfeld, Ifat Kaplan-Ashiri, Ori Avinoam, Chen Speter, Moshe Halak, Howard Kruth, Lia Addadi

**Affiliations:** Department of Chemical and Structural Biology, Weizmann Institute of Science; Rehovot, Israel 7610001; Department of Chemical Research Support, Weizmann Institute of Science; Rehovot, Israel 7610001; Department of Biomolecular Sciences, Weizmann Institute of Science; Rehovot, Israel 7610001; Department of Vascular Surgery, Sheba Tel HaShomer Medical Center; Ramat-Gan, Israel 52621; Experimental Atherosclerosis Section, National Heart, Lung, and Blood Institute, National Institutes of Health; Bethesda, Maryland, 20892

## Abstract

Atherosclerosis is a pathology affecting the arteries, characterized by the buildup of plaques in the blood vessel walls. Atherosclerosis is the main cause of cardiovascular diseases, which constitute the leading cause of death in the world. Cholesterol crystals are the main components of the plaques, which actively participate in plaque growth and rupture and do not dissolve in aqueous environments. Employing novel cryo-scanning electron microscopy techniques, we examined human atherosclerotic plaques at high resolution, in 3D, and in close to native conditions. We show that cholesterol crystal clearance occurs in advanced human plaques through the activity of cells. We suggest that this occurs by enzymatic esterification of cholesterol to cholesteryl ester, which aggregates into intra- and extra-cellular pools. This discovery provides further understanding of the disease process in atherosclerosis, and may inspire new therapeutic approaches.

## Introduction

Atherosclerosis is the main reason for heart attacks and strokes, which constitute the leading cause of death worldwide (*1, 2*). Atherosclerosis is a chronic vascular disease that initiates at a very young age with the formation of plaques in the artery wall, composed mainly of lipids, cell debris, cholesterol crystals and, later, calcifications (*3-5*). Cholesterol crystals are crucial in the clinical manifestations of atherosclerosis due to their detrimental effects on plaque stability and rupture (*6-8*). Cholesterol crystals are present even in the early stages of lesion formation and are commonly associated with macrophages and smooth muscle cells, major cellular players in the developing lesion (*3, 9-11*). Cholesterol crystallization is considered a unidirectional process in the human lesion, because cholesterol crystals are insoluble in aqueous environments (*3, 12*). In several animal models, cholesterol and other lipid depletion occurred in atherosclerotic lesions after pharmacological interventions or after prolonged periods during which cholesterol levels in the blood were kept normal (*13-16*). Cholesterol crystal dissolution was never directly documented in human plaques, although Abela’s group reported that treatment by incubation of excised atherosclerotic tissues with statins or aspirin showed a decrease in plaque volume and dissolution of exposed crystals (*17, 18*).

Employing high-resolution cryo-scanning electron microscopy imaging techniques, we examined human atherosclerotic lesions in the hydrated state, close to native conditions. We show evidence for the role that intralesional cells play in cholesterol crystal breakup, even in very advanced stages of atherosclerotic lesions.

The cryo-conditions that we used allowed us to observe cholesterol deposits and crystals in minimally processed diseased tissue. We believe that intralesional cell-mediated cholesterol crystal breakup in human atherosclerotic plaques was not previously discernable because most high-resolution imaging techniques use embedding, which entails the dissolution of all lipid components from the tissue during sample preparation (*19-22*). The studies using scanning electron microscopy involved, as a minimum, drying of the sample, which may cause shrinking and deformation of the tissue, evident especially at high resolution (*23-25*).

## Results

### Imaging hydrated human atherosclerotic tissues in 3D, close to native conditions

Twelve human atherosclerotic tissues were excised via an elective endarterectomy of the carotid artery (*26, 27*). The tissues excised are for the most part very voluminous (∼4-5 cm long, lesions 0.020-5mm thick), although they comprise only the diseased intima region of the arterial wall (Fig. S1). They often contain macroscopically evident deformations and multiple local atherosclerotic centers, some of which are necrotic. The excised plaques (Fig. 1A, SI 1) are generally rich in fibrous components and hard calcification deposits throughout (Fig. 1A, B features in white).

**Fig. 1.**
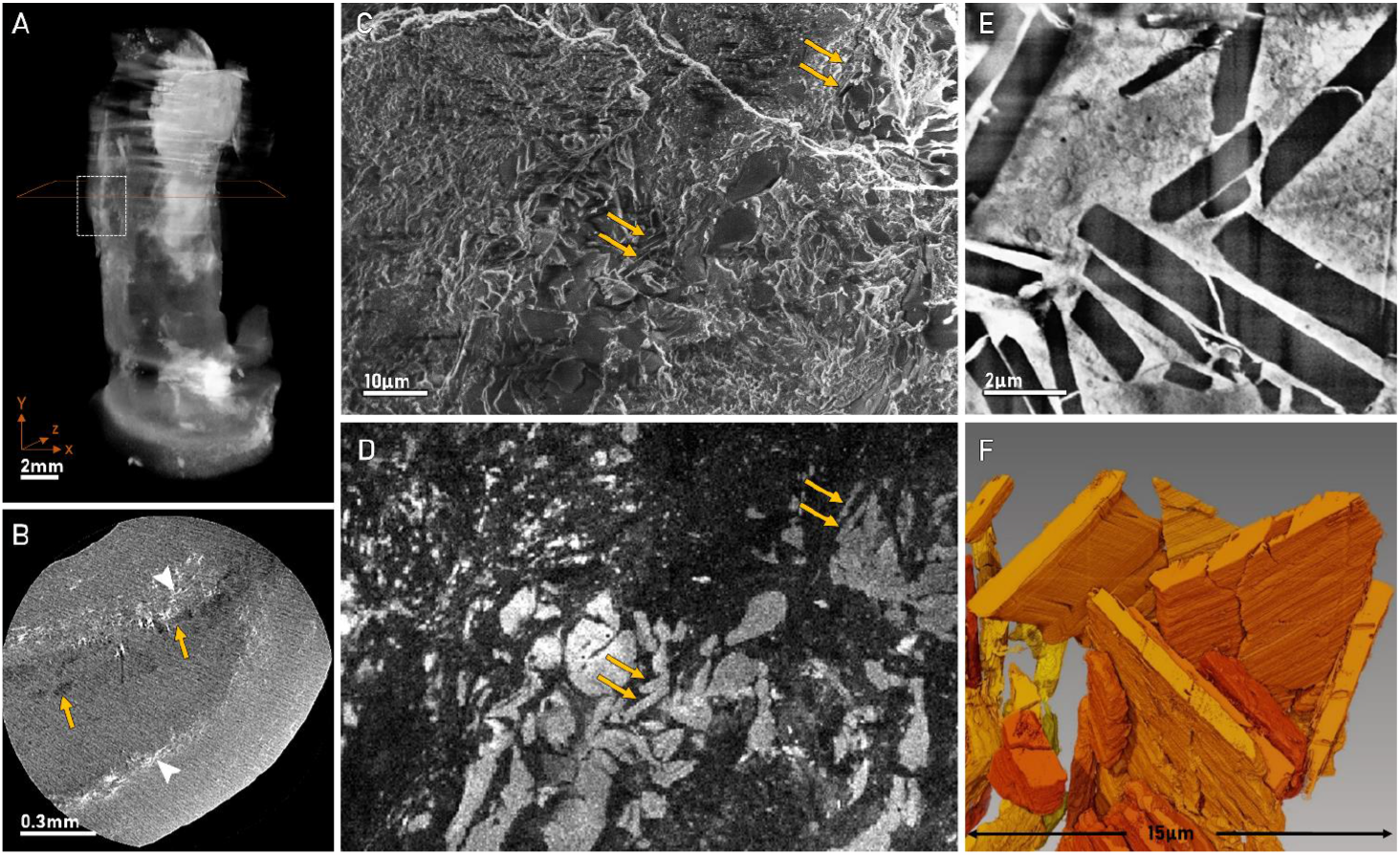
Imaging human atherosclerotic lesions cm to nm scale - workflow procedure. (samples taken from patient #3) **A-B**) Micro-CT. A) Volume reconstruction of the whole lesion (movie S1). The voxel size is ∼22.7 µm; B) Individual slice, the voxel size is ∼2.71 µm. Cholesterol deposits (yellow arrows) are dark grey whereas calcifications (white arrowheads) are white. **C-D**) Cryo-SEM of high pressure frozen and freeze fractured sample. C) Secondary electrons image: individual crystals are difficult to detect in the cell debris and lipid-rich environment; D) cathode luminescence (CL) image of the same frame as in C): individual cholesterol crystals are detected by virtue of their bright white luminescence (e.g., crystals labeled with yellow arrows in C and D). Calcifications do not have appreciable signal in cryo-CL in the experimental conditions used here. **E–F**) Cryo-FIB-SEM. E) Individual slice, showing an aggregate of large crystals (black). Note that in micro-CT and in cryo-FIB-SEM cholesterol deposits are black, whereas in CL images, cholesterol deposits appear white. F) Surface representation of the crystal aggregate in E after 3D image reconstruction and segmentation.

A workflow procedure starting from low-resolution-high-volume imaging and ending with high-resolution-low-volume imaging was established, to allow data acquisition in a wide range, from cm to nm scale (Fig. 1). Micro-CT was first performed to acquire information on the inner structure, components and lesion locations, encompassing the whole excised volume (Fig. 1A, B and movie S1)(*28*). Differences in the density of the tissue components allowed detection of the position and size of calcifications (bright white) and of large cholesterol crystal stacks (dark grey, Fig. 1B) in the lesion tissue (grey). Most of the tissues exhibited multiple large lesions with heterogeneous composition.

Based on the picture provided by micro-CT, we selected appropriate regions of interest, in the core and in the periphery of lesions, as candidates for inspection at higher resolution. In general, we tried to avoid regions with extended calcifications, concentrating on regions that contained cholesterol deposits. The selected regions were dissected, high pressure frozen (HPF) to produce hydrated vitrified samples, and freeze fractured. The frozen freeze fractured samples were imaged by cryo-scanning electron microscopy (cryo-SEM) (Fig. 1C) equipped with cathodoluminescence (CL) detection capability (Fig. 1D). The combination of cryo-SEM with CL allows identification of cholesterol and cholesteryl ester deposits, due to their luminescent properties upon excitation with an electron beam (Fig. 1C, D)(*29*). Lesion location/boundaries and correct identification of cholesterol interaction with the tissue components are otherwise difficult, because of the loss of architecture of the diseased tissue.

Finally, localized regions of interest were selected and imaged in 3D with cryo-Focused Ion Beam SEM (cryo-FIB-SEM) (Fig. 1 E, F) (*29, 30*).

A large burden of cholesterol crystals arranged in stacks or as individual crystals characterized the core regions of the lesions, and is presented in a slice from a cryo-FIB-SEM stack in E, followed by a volume representation of the crystals after image segmentation in F.

### Cholesterol crystal breakup within the atherosclerotic lesions

Examination of the lesion tissues with a combination of cryo-SEM and cryo-FIB-SEM led to the detection of strong evidence for cholesterol crystal breakup and disintegration in the human lesions. Fractured crystals display jagged edges that change shape and regress in subsequent sections from cryo-FIB-SEM stacks, showing patterns typical of dissolving crystals (Fig. 2A, B, compare to Fig. 1E) (*31*). A multitude of round water-filled vesicles surrounds the areas where the crystals appear fragmented and intimately associated with the crystals. Often, although not always, we can observe cellular membranes enveloping the mass of globular bodies/vesicles together with the fragmented crystal area, creating an isolated volume where controlled processing may occur (Fig. 2A). In cryo-SEM images as well, jagged crystals appear completely surrounded by globular bodies of sizes 0.1-1 µm (Fig. 2C, D). Some of the globular bodies are located inside cavities in the crystals (Fig. 2C, inset). The cavities match the globular bodies in size and shape, suggesting that the globular bodies themselves may be actively involved in the breakup of otherwise smooth crystals (Fig. S2 A, B).

**Fig. 2.**
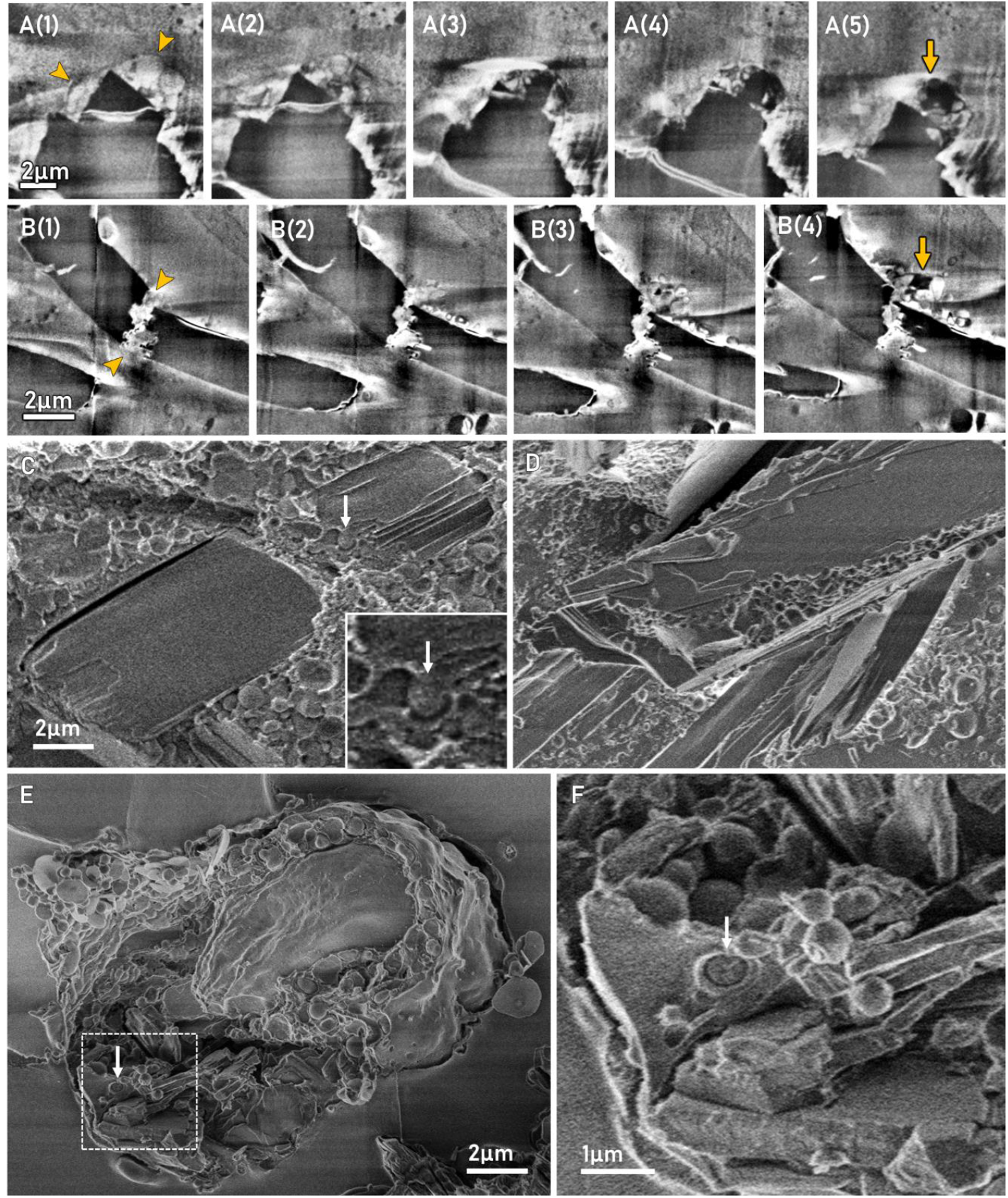
Cholesterol crystal breakup. **A-B**) Subsequent sections from cryo-FIB-SEM stacks carved in an atherosclerotic lesion extracted from patient #3. A1-A5) A triangular chip breaks off from a crystal (black). The chipped fragment is surrounded by a membrane (yellow arrowheads in A1), delimiting a space occupied by white water-filled vesicles and a black cholesterol deposit (yellow arrow in A5). B1-B4) The crystal (black) has jagged edges (yellow arrowheads in B1). White water-filled vesicles and black cholesterol deposits (yellow arrow in B4) surround the jagged edges. In cryo-FIB-SEM, where contrast depends on surface potential, cholesterol and cell membranes appear dark, whereas bright areas indicate polar aqueous solutions. **C-D**) Cryo-SEM images of cholesterol crystals from an atherosclerotic lesion extracted from patients #4 and #3, respectively. Globular bodies surround the crystal, some penetrated inside the crystal (arrow in C, magnified in the inset). **E-F**) Cryo-SEM images of a J774A.1 macrophage cultured for 12 hours in the presence of cholesterol crystals. Globular bodies surround several phagocytosed crystals, some penetrated inside the crystal (arrow in F, magnified from the white frame in E).

Further support for this hypothesis derives from cryo-SEM observation of J774A.1 macrophage cultures (controls in Fig. S2 C, D) exposed to crystals of cholesterol (crystals shown in Fig. S2 A, B). After 12 hours, the cells actively phagocytosed large amounts of cholesterol crystals. In contrast to the initial crystals the phagocytosed crystals have jagged borders, and many globular bodies populate the cell cytosol (Fig. 2E). Similar to what we observed in the crystals within the atherosclerotic lesion, some of the globular bodies appear to penetrate into the crystals (Fig. 2F).

### Cellular involvement in cholesterol crystal processing and breakup

The observations so far mentioned in the human lesions were made in cellular regions, identified by the presence of cell nuclei and organelles, suggesting intra-cellular locations, although whole cell membranes could not always be delineated (Fig. 3A). In the cellular region imaged by cryo-FIB-SEM in Fig. 3A) and movie S2, we observed a membrane-delimited space, which completely encompasses a mature crystal, and contains smaller vesicles/globular bodies. Large (3-10 µm diameter) ‘foamy structures’ appear (Fig. 3A, yellow arrowheads), often intimately associated with crystals (Fig. 3B), composed of dark and light areas, mostly bright vesicles on a black background. Similar foamy structures were detected in all the tissues of patients analyzed with cryo-FIB/SEM. We note that in cryo-FIB-SEM, contrast originates from surface potential and is dependent upon the interaction of biological materials with the electron beam. Hydrophobic materials in general, and specifically cholesterol and cell membranes, appear dark because they accept and retain electrons, whereas bright areas indicate in this context polar aqueous solutions that repel and emit electrons (*30, 32*).

**Fig. 3.**
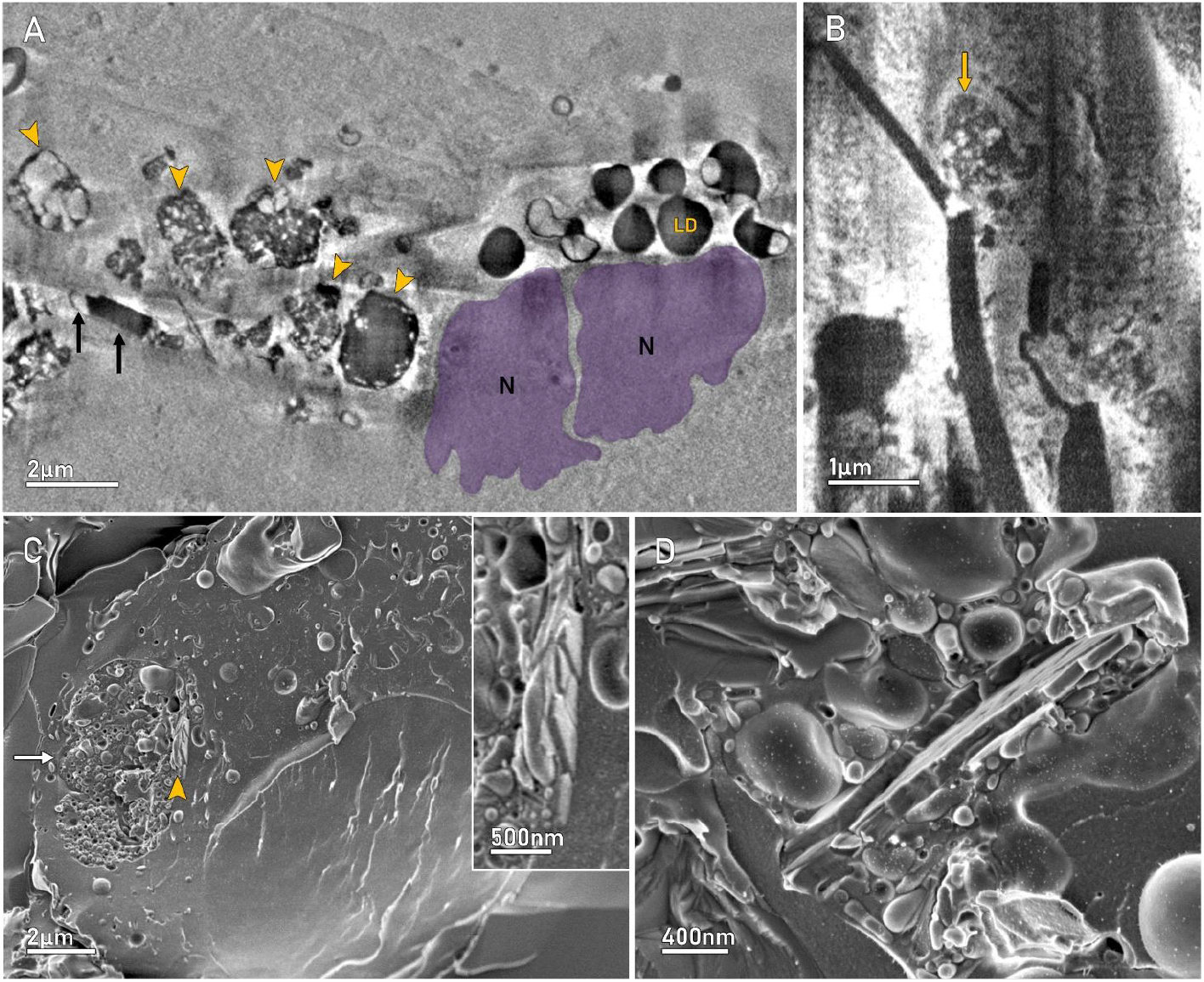
Cellular involvement in cholesterol crystal processing and breakup. **A-B)** Cryo-FIB-SEM slices carved in atherosclerotic lesions excised from patient #3 and patient #7, respectively. A) One membrane-bound crystal (black arrows), several lipid droplets (black) and several foamy structures (arrowheads) presenting white (water-filled) vesicles on a black cholesteryl ester background, in a cellular environment. The cells are identified by the presence of intact cell nuclei (purple, N), and intracellular lipid droplets (LD). B) Foamy structures at a site of crystal dismantlement (arrow). **C, D)** Cryo-SEM images from a J77A4.1 macrophage cultured with cholesterol crystals for 4 hours, followed by removal of the crystals and incubation for additional 20 hrs. C) Note the large foamy structure (arrow) and the remainders of crystals intimately associated with it (arrowhead, magnified in the inset). D) Crystal enclosed in a foamy structure, containing also larger droplets.

In macrophage cultures exposed to cholesterol crystals for a relatively short time of four hours, we observed, using cryo-SEM, crystals immersed in liquid contained inside membrane-limited bodies mixed with complex organelles, clearly water-filled (Fig S3). Also, large dense non-membrane-limited bodies appear (yellow arrows in Fig S3), which we infer correspond to the dark hydrophobic material in cryo-FIB-SEM and are lipid droplets (LD in Fig 3A).

After further incubation for up to 24 h of the macrophage cultures exposed to crystals for 4 h, the few crystals left inside the cells display characteristics of dissolution (Fig 3C, inset). Some of the crystals are associated with large bodies containing many small vesicles (Figure 3C, D), which we identify with the ‘foamy structures’ observed in the atherosclerotic lesions (Fig 3A, B)

The extracellular regions of the lesions are rich in cholesteryl esters forming extended pools (*3, 33*). Some of these cholesteryl ester pools, observed in cryo-SEM, display the typical pattern of cholesteric liquid crystals of cholesteryl esters (Fig 4A and SI4), as it appears in the literature for artificial samples of cholesteryl esters (*34*), allowing their identification as such. Most poly-unsaturated cholesteryl esters at ambient or even higher temperatures, pack as liquid crystals. Based on this structural packing behavior, Small suggested, although he could not demonstrate it at the time, that large cholesteryl ester pools in atherosclerotic lesions might assume a liquid crystalline phase (*3, 35-37*).

**Fig. 4.**
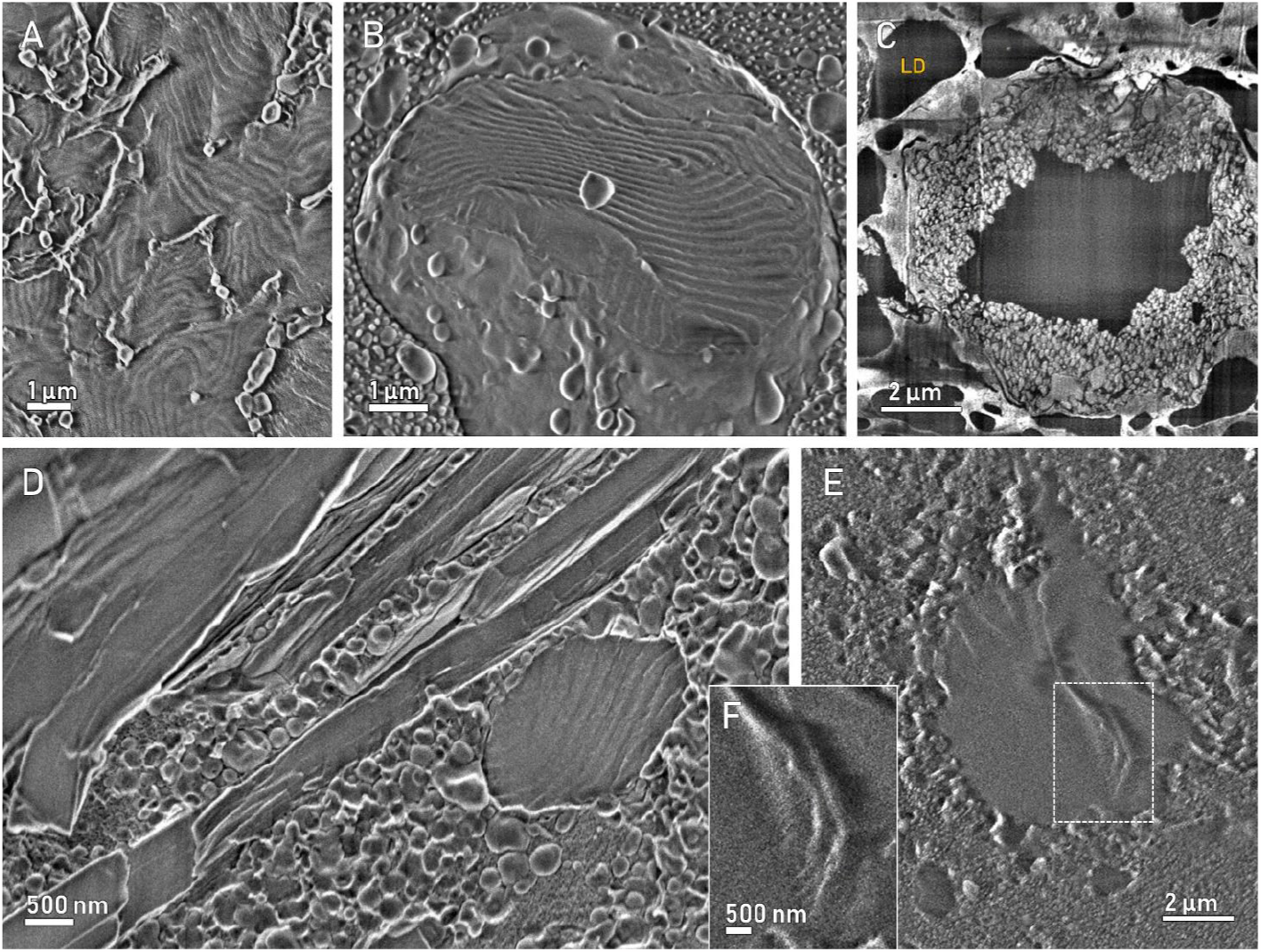
Cholesteryl ester pools in lesion tissues. **A)** Cryo-SEM micrograph of extracellular cholesteric liquid crystalline pools of cholesteryl esters. **C)** cryo-FIB-SEM slice of a foamy structure containing a non-faceted hydrophobic core. **B, D, E)** Cryo-SEM images of foamy structures as in C), from patients #3, #4 and #5. The hydrophobic cores are identified as cholesteryl ester by the characteristic cholesteric liquid crystal patterns. The inset **F** is a magnification of the boxed area in E, better showing the liquid crystal pattern. The texture and the spacing of the periodic structures, 140-180nm, are identical to those of extracellular liquid crystalline pools as in A), and are within the range of the periodic spacing of cholesteryl ester liquid crystals (*34*).

We observed in cryo-FIB-SEM of lesion tissues, relatively large cellular foamy structures comprising a dark non-crystalline core and a vesicle-packed shell (Fig 4C). In cryo-SEM, some of these same structures, often observed in contact or in the immediate vicinity of crystals, displayed in their core the same cholesteric liquid crystal patterns (Fig 4B, D, E) as the extracellular cholesteryl ester pools, thus disclosing that their composition is also cholesteryl ester.

## Discussion

Considering that cholesterol crystals are practically insoluble in water environments, what can be a possible mechanism for cholesterol crystal breakup within an advanced lesion? The relatively large foamy structures observed in the lesions, comprising a cholesterol-containing core and a vesicle packed shell appear to be filled with cholesteryl esters, because they display the characteristic texture of cholesteric liquid crystals. Their locations and their appearance support the notion that the non-vesicular content of the foamy structures is cholesteryl ester that results from cellular activity on the cholesterol crystals (*38*). We suggest that crystal processing and subsequent breakup occur through cell-mediated esterification of the cholesterol molecules to cholesteryl ester.

The aqueous content of the vesicles found in the periphery of the structures, together with the evidence of active contribution to crystal resorption by vesicles burrowing inside the crystals, further support the notion that breakup is carried out through an enzymatic activity in aqueous compartments (*33*). This implies that crystal breakup is achieved through chemical transformation, rather than by dissolution in an aqueous environment. The chemical transformation most probably occurs via transposition of cholesterol molecules from the cholesterol crystals to the vesicle hydrophobic membrane, and subsequent exposure to esterification in the intra-vesicular aqueous environment (*9, 39*). The newly formed esters can then self-assemble to form the dark regions in the foamy structures associated with crystals (Fig. 2A, B, Fig 3 A, B) and the large liquid-crystal cores (Fig. 4 B, D, F). Published TEM images bear resemblance to the ‘foamy’ structures that we have observed, some of them referred to as auto-lysosomes (*38, 40*).

The process of esterification can occur by the reverse enzymatic function of neutral and/or lysosomal esterases, and ACAT, the enzyme that performs esterification of cholesterol. Supporting this hypothesis, Sakashita et al reported that upon supplementation of human macrophages with cholesterol, the cells produce vesicles enriched with ACAT (*41-43*). These vesicles, which originate in the ER and co-localize with late endosomes/lysosomes, strongly resemble the vesicles that we observe in the shell region of the foamy structures (*42*). Furthermore, cholesterol esterases can catalyze both the forward and backward reactions of cholesterol hydrolysis and esterification, respectively. Specifically, hydrolysis is preferred at the earlier stages of the disease, but esterification is favored at the later stages (*44-47*). McConathy et al, Rajamäki et al and Kruth et al showed that macrophages phagocytose and process cholesterol crystals transforming the cholesterol molecules into cholesteryl ester, detected by analytical methods in intracellular cholesteryl ester pools (*38, 48, 49*). While evidence for crystal processing by cholesterol esterification has been put forward in macrophage cultures, no indication of cells being involved in the process of cholesterol crystal clearance in human atherosclerotic lesions existed until now.

Macrophages and/or other cells in atherosclerosis are conventionally considered associated with cholesterol crystallization, rather than breakup (*11, 48-51*), especially in the developed and mature lesions, where compensatory mechanisms for the inhibition of cholesterol crystallization are exhausted (*4, 21*). Most atherosclerotic lesion studies were performed on animal models. In these experiments, the animals are artificially administered a high cholesterol diet for a relatively short period, to induce rapid atherosclerotic lesion development (*15, 52*), whereas human atherosclerotic lesions develop and mature for decades. For this reason, animal lesions are under active construction when observed, and new cholesterol crystal deposition occurs, rather than crystal breakup. Crystal clearance, as demonstrated here, is more pertinent to mature lesions, as those excised from humans.

Cholesterol crystals in atherosclerotic lesions are dangerous for several reasons: they increase the plaque probability to fracture, generating emboli that trigger major cardio-vascular events, they cause inflammation and cell death, and they are very difficult to dissolve in an aqueous environment. Here we show that inside advanced human atherosclerotic plaques intralesional cells are active in dismantling cholesterol crystals. We observe the intimate association between crystals, vesicles with aqueous content and cholesteryl ester-containing bodies that populate the intra- and extra-cellular space. We infer from our observations that cholesterol crystal breakup in the advanced human lesion proceeds through cholesterol esterification to cholesteryl ester. Cholesteryl esters are physiological cell components, do not have mechanical impact, and can be transported in the blood. Cholesterol crystal clearance by macrophages or other intralesional cells may thus constitute a mechanism of self-defense of the body, opening the way to potential therapeutic approaches to the threats inherent to atherosclerosis and other diseases.

## Materials and Methods

### Cholesterol Monohydrate Crystal preparation

Cholesterol crystals were prepared by crystallization of powdered cholesterol (Avanti Polar Lipids Inc.) from lab-grade acetone (Bio Lab ltd., Israel). 62.5 mg cholesterol powder was dissolved in 5 ml acetone by heating. 20 ml water was added to the solution and the mixture was let cool 20-30 min while stirring. The crystals were sonicated to reduce clumping and the solution was filtered through a sinter glass Buchner funnel filter (grade 3), washed three times with DDW and dried. Successful crystallization was confirmed by polarized light microscopy and by SEM (Zeiss Sigma 500, Oberkochen, Germany) of crystals coated with 5 nm of Iridium (Safematic CC-010 HV). Crystal size is 2-5 µm. Prior to the use in experiments, the crystals were sterilized under UV for 1 hour.

### Cell Culture

Murine macrophage-like J774A.1 cells were provided by the lab of Prof. Irit Sagi, Department of Immunology and Regenerative Biology, Weizmann Institute of Science, Israel. Cells were cultured in DMEM (Dulbecco’s Modified Eagle Medium, Gibco), supplemented with 10% fetal bovine serum (FBS, Sigma-Aldrich #F7524), 1% L-Glutamine, 100 units/ml penicillin and 100 g/ml streptomycin (Pen-Strep solution, Sartorius #03-031-1B). Cultures were maintained at 37ºC and 5% CO_2_. Samples were prepared by seeding 3.5 or 5 million cells on 0.04 g of sterile synthetic cholesterol crystals in a 100 mm Falcon cell culture dish (Corning #353003). The dishes were incubated for 4 or 12 hours. After incubation, the dishes were washed in warm DPBS (Dulbecco’s phosphate buffered saline, Sartorius, #02-020-1A) to remove floating crystals. In the pulse-chase experiments, the cells were incubated with the crystals for 4 hours before washing, and for further 20 hours after washing. The cells were scraped and centrifuged for 5 minutes at 180 g before high pressure freezing (described below).

### Human atherosclerotic lesions

Use of all human tissues strictly adhered to accepted guidelines of the Helsinki declaration and was made upon approval of the Helsinki committee of Sheba Tel HaShomer Hospital, Israel and the approval of the Institutional Review Board of the Weizmann Institute of Science, Israel. Twelve human atherosclerotic tissues ranging from 3-5 cm in length and 1-1.5 cm in diameter, were excised via an elective endarterectomy procedure. Informed consent was obtained from all patients after the nature and possible consequences of the studies were explained. Upon excision, the tissues were immediately placed in 0.1 M cacodylate buffer, containing 5 mM CaCl_2_ and 4% paraformaldehyde (PFA) at pH 7.4. Following 24 hours, tissues were placed in fresh 0.1 M cacodylate buffer, containing 5 mM CaCl_2_, 4% PFA and 0.5% glutaraldehyde (GA) at pH 7.4 and stored at room temperature until further processing.

### Micro-Computed Tomography (Micro CT)

Micro CT scans were obtained using a Zeiss Xradia Versa 520 (Zeiss X-ray Microscopy, California, USA). Atherosclerotic tissues from 10 patients were washed in DDW and placed in a designated tube. 0.2 ml DDW was placed at the bottom of the tube to allow tissue hydration throughout the duration of the scan, the tube was sealed with a rubber cap to prevent water evaporation. The portion of the tissue to be scanned was not immersed in water but was completely exposed to air in a hydrated environment. These settings allowed maintaining a sufficient contrast and resolution while maintaining tissue hydration. 1600 projections were executed over 360°, at 30 kV and 2 W. Scans were performed at voxel size of 2.71 µm^3^, 9.73 µm^3^ or 22.71 µm^3^. Volume reconstruction was performed with Zeiss proprietary software based on the back projection algorithms.

### Cryo-Fixation

Human atherosclerotic tissues were examined using a ZEISS Stemi 305 Compact Greenough Stereo Microscope and underwent slicing according to the regions of interest chosen based on the micro-CT data. Tissues were sliced to a thickness of 100-200 µm and 0.5-1 mm width. Macrophage cells were gently scraped from the culture dishes and centrifuged (180 g, 5 minutes) to form a concentrated cell pellet. Samples were sandwiched between two aluminum discs of 3 mm-diameter, 0.025-0.1 mm depth (Engineering Office M. Wohlwend GmbH, Switzerland), while immersed in 0.22 µm pore-sized syringe filtered PBS with 10% dextran (Sigma-Aldrich, #31389). Samples were cryo-immobilized using a high-pressure freezing device Leica EM ICE (Leica microsystems, Germany). The sandwiched frozen samples were stored in liquid nitrogen until freeze fracturing and examination. For freeze fracture, a sample was transferred into a freeze fracture device (BAF 60, Bal-Tec; ACE900, Leica) employing a Vacuum Cryo-Transfer unit (VCT) 100 (Leica Microsystems, Wetzlar, Germany) while maintaining vacuum conditions of 6x10^-7^ mbar and cryogenic conditions. Finally, the sample was transferred using the VCT 100 to a Gemini SEM 500 or Ultra 55 SEM (Zeiss, Oberkochen, Germany) where it was inspected at -120°C, or to a Cross Beam (Zeiss, Oberkochen, Germany) where it was maintained at -150°C for cryo-FIB-SEM milling and imaging. Macrophage samples that were imaged in the Gemini SEM 500 were also etched for 1 minute at 110°C and coated with 6 nm of Carbon at 165 W in the freeze fracturing device.

### Cryo-SEM

Tissues were studied in hydrated vitrified conditions (*29, 50*). Vitrified samples were mounted on a holder under liquid N_2_, transferred to a BAF 60 device and fractured at -120°C, 6 x10^-7^ mbar. Samples were then shuttled to a Zeiss Gemini SEM 500 at cryogenic conditions at 6 x10^-7^ mbar, were mounted on a cryo-stage and etched for 15 min at -105°C. Tissues were imaged at 1-2 kV, -120° C.

### Cathodoluminescence (CL)

The CL detector and spectrometer is installed on a Zeiss Gemini SEM 500 and is comprised of a Gatan MonoCL4 Elite system equipped with a retractable diamond-turned mirror. The collected light was recorded using a photomultiplier tube in panchromatic mode. CL measurements were performed using a SEM aperture of 15-30 µm, low exposure times (150 µs/pixel), at 4-5 kV. The temperature was maintained between -120 and -140°C. Sample preparation was as for the cryo-SEM imaging.

### Cryo-FIB-SEM

The method provides good 3D imaging of relatively large volumes of tissues at a resolution of up to tens of nanometers (*29, 30*). The technique combines serial focused ion beam milling of cryo-fixed (vitrified) samples with sequential SEM block face imaging under cryogenic conditions. Acquisition of serial images is achieved over large volumes of tens of micrometers cubed, at a voxel size of up to 5 nm^3^. Sample preparation for cryo-FIB-SEM does not require chemical fixation and/or the use of contrasting agents. Vitrification of samples was performed by HPF, as described above. A vitrified sample was transferred to a Vacuum Cryo Manipulation loading station and the frozen aluminum disc was mounted on a cryo-holder under liquid N_2_. The sample was then transferred to a BAF 60 device using a VCT 100, where it underwent a fracture at -150°C and coating with a 6-10 nm-platinum layer to increase sample conductivity and reduce charging effects. Samples were then transferred using a VCT 100 to the Gemini SEM 500, equipped with a CL detecting system, where a CL map was recorded. The sample was then transported using the VCT 100 unit to the Crossbeam and the region of interest was located using the collected CL map data. The cryo-stage was tilted at 54° and stage height was adjusted to a working distance of 5 mm. An organometallic protective layer was deposited on top of the fractured surface using a gas injection system (GIS) heated to 27° C. The working distance during deposition was increased to 7.5 mm. The deposition process consisted of opening the outlet valve of the GIS system for 30 seconds, twice, which results in condensation of organometallic composition on the sample with a layer of about 100-300 nm.

Trenches of approximately 90X40 µm were milled just in front of the area of interest, and data collection was performed consecutively in the designated positions as described below. 2-4 stacks were collected from regions proximal to the lesion cores of 5 different patients. Slices number for each stack ranged within 879-2507, with an average slice area of 30x40 µm^2^ and a constant slice thickness of 20 nm for all stacks. The following milling and imaging parameters were applied for the data-set acquisition: for the initial trenches 30 kV, 7-15 nA were employed. 1.5-3 nA with a dose factor of 40-50. For the two-stack data presented in this manuscript, the following are the specific volume parameters: Stack 1: 1842 slices comprising a total volume of 40x30x36.8µm^3^. Stack 2: 1487 slices comprising a total volume of 40x30x29.7 µm^3^ SEM images were recorded at 2.2 kV, 50 pA with a pixel size of 19.53 nm.

### Image analysis

All 2D images acquired via Cryo-SEM were subjected to level adjustment using Adobe Photoshop software (Adobe Photoshop CS4 Extended, Middle Eastern Version 11.0).

For data analysis acquired with Micro CT, volume generation was performed, and image contrast was adjusted, using Avizo Lite 9.2.0 software.

For data analysis acquired with cryo-FIB/SEM, elimination of periodic non distorted vertical and horizontal linear artifacts was made by the FFT bandpass filter tool in Fiji/ Image J (National Institute of Health, Maryland, USA). To allow better detail detection of intra/intercellular structures, local contrast enhancement (CLAHE) tool in Fiji/ Image J and an “unsharp masking” filter in Avizo or Fiji/Image J were applied. The images were aligned using Fiji “Linear Stack Alignment with SIFT” plugin, and a more refined alignment was performed manually using the alignment tool in the Avizo Lite 9.2.0 software. Segmentation: Manual segmentation of features of interest was performed. Segmented features were displayed via volume generation or surface generation, with 3–4-unit smoothing effect. We note that the noticeable periodic striation on the segmented crystal surfaces along the milling direction are artificial and derive from the data acquisition and segmentation processes.

## Acknowledgments

We thank Dr. Neta Varsano for her help with cryo-SEM and image processing. We thank Leslie Leiserowitz and Benjamin Geiger for critical reading of the manuscript. We thank Prof. Irit Sagi for the gift of J774A.1 cells.

