## Supplementary Information for "Cell-mediated cholesterol crystal processing and clearance observed by 3D cryo-imaging in human atherosclerotic plaques"

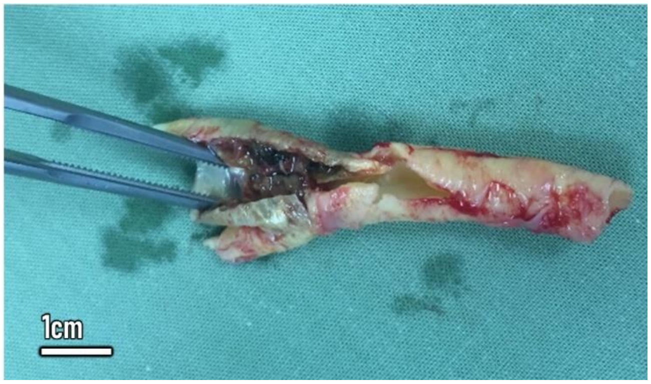


**Fig. S1.**
Human atherosclerotic tissue excised from a carotid artery via an endarterectomy procedure.

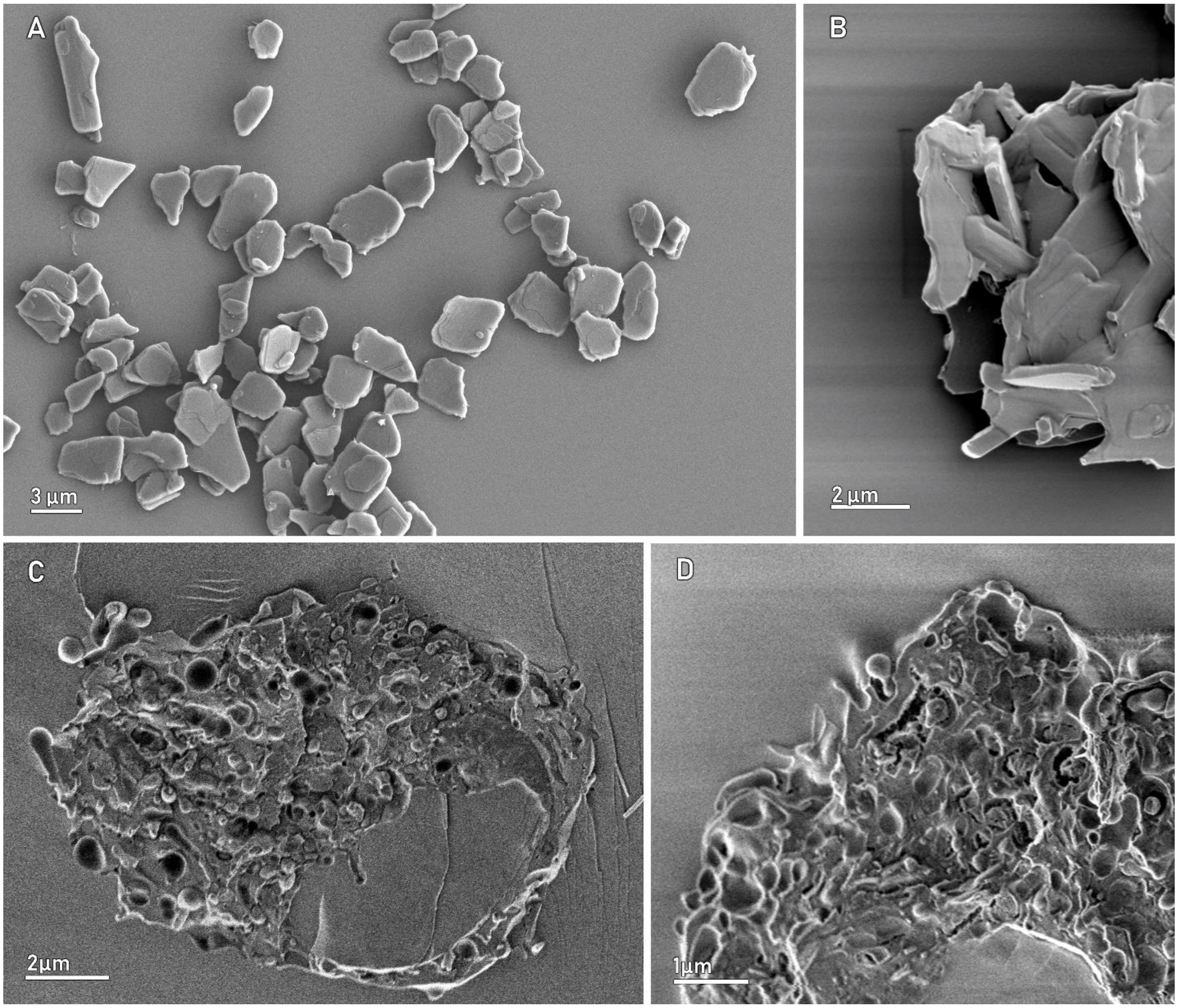


**Fig. S2.**

**Controls**. **A, B**) Cholesterol crystals from the same batch as those incubated with the cells. **C, D**) Cryo-SEM images of J77A4.1 macrophages cultured under the same conditions as those in Fig. 2-3, without crystals.

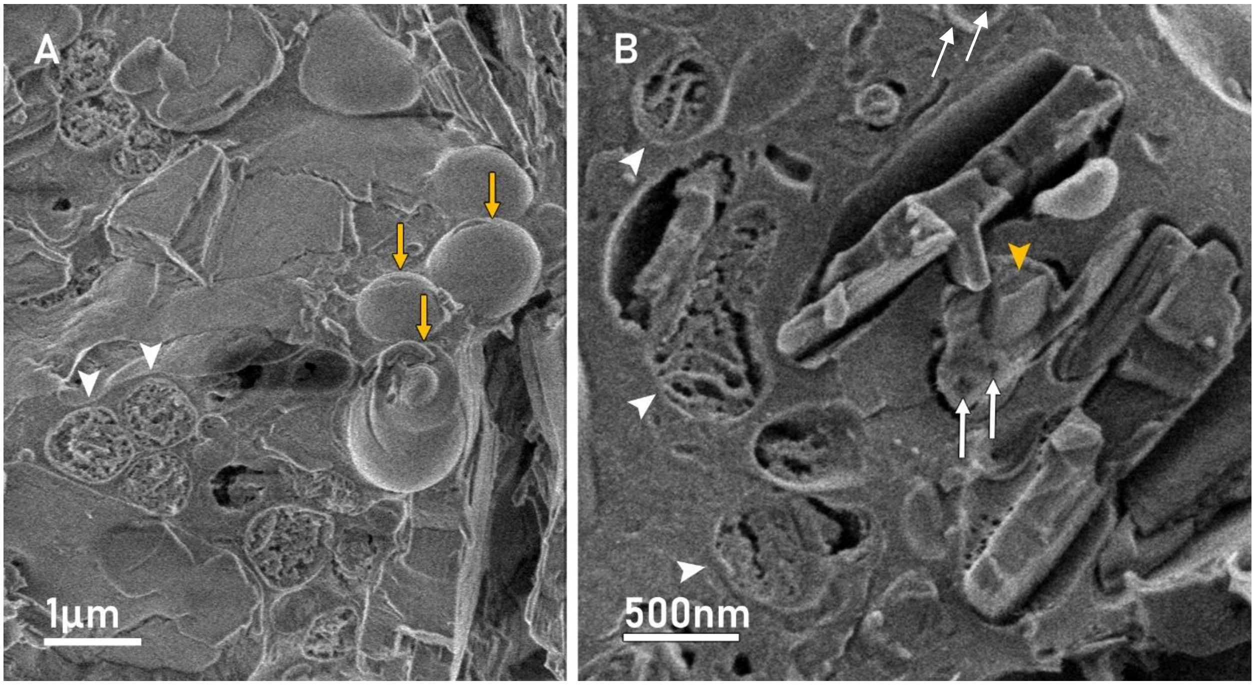


**Fig. S3.**
Cryo-SEM images from J77A4.1 macrophages cultured with cholesterol crystals for 4 hours. **A)** Yellow arrows indicate lipid droplets, whereas white arrowheads indicate water-containing organelles with complex microstructures, most probably endosomes or lysosomes. These organelles are very abundant in the cells incubated with crystals. **B)** Crystals are inside membranes (yellow arrowhead) that may contain dense material and distinct vesicles (white arrows). Different water-containing organelles with complex microstructures, possibly endosomes or lysosomes, are abundant (white arrowheads).

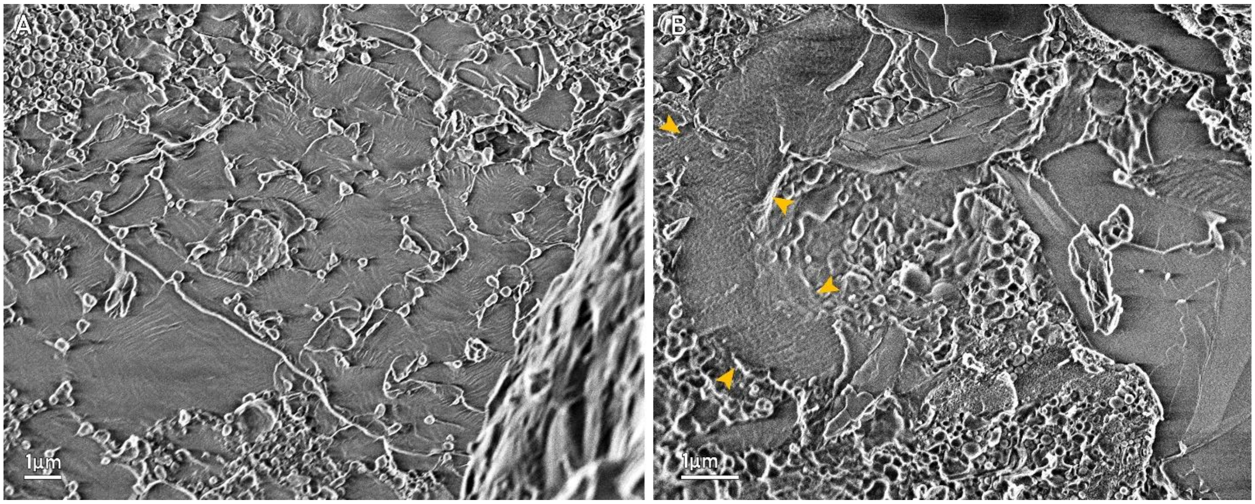


**Fig. S4.**

Cryo-SEM micrographs of extra-cellular cholesteryl-ester liquid crystalline pools. The yellow arrowheads indicate regions where the liquid crystal patterns are evident.

**Movie S1.**

https://weizmann.app.box.com/s/x6f52bhidkob2goxnkpthwc8ypivt22z

**Micro CT volume representation of an intact excised atherosclerotic tissue.** Higher resolution of the tissue was acquired and registered in the complete volume, followed by segmentation: soft tissue (light grey), lipid core (orange) and calcium deposits (white). Subsequent transverse slicing of the tissue volume: soft tissue (grey), lipid core (dark grey), calcium deposits (white).

**Movie S2.**

https://weizmann.app.box.com/s/mld146h1ruh33xm5cwd7r8zv9mz41sfa

**Consecutive cryo-FIB-SEM slices from a cellular region in the atherosclerotic lesion**. 1062 slices, slice thickness = 20 nm. Field of view is 22.3X17.95 $\mu m^2$. **A**) Individual snapshot showing cell nuclei (N) juxtaposed to foamy structures (black arrows). **B**) Individual snapshot where the distinction between lipid droplets (LD) and foamy structures (black arrows) can be observed, plasma membrane is indicated by red arrowheads.

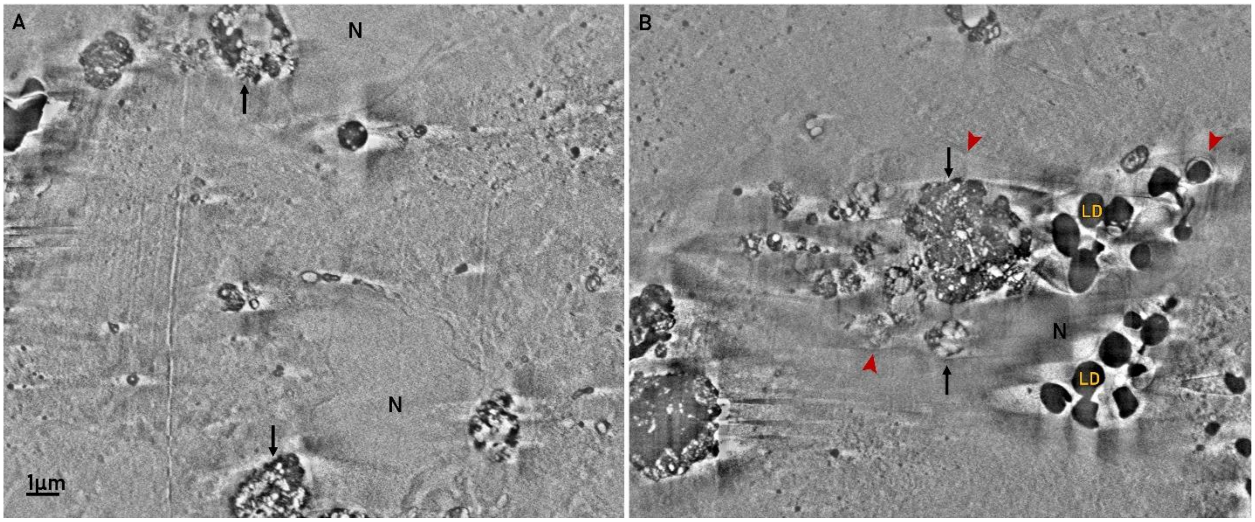
